## Appendix for "Structured Peer Review: Pilot results from 23 Elsevier Journals"

This is the appendix for ***Structured Peer Review: Pilot results from 23 Elsevier Journals*** by Mario Malički and Bahar Mehmani. The order of items is based on how the information is presented in the manuscript.

### Appendix Table 1. Journals per field and impact factor quartile.

| **Field** | **IF Quartile 0** | **IF Quartile 1** | **IF Quartile 2** | **IF Quartile 3** | **Total** |
| --- | --- | --- | --- | --- | --- |
| Health and Medical Sciences | 1 | 2 | 1 | 1 | 5 |
| Life Sciences | 2 | 2 | 2 | 1 | 7 |
| Physical Sciences | 0 | 2 | 2 | 2 | 6 |
| Social Sciences and Economics | 0 | 1 | 2 | 2 | 5 |
| Total | 3 | 7 | 7 | 6 | 23 |

We analysed 107 manuscripts across 23 journals (20 journals had 5 manuscripts with 2 reviews, 1 journal had 4 such manuscripts, 1 had 2 such manuscripts, and the final 1 journal had only 1 such manuscript).

### Appendix Box 1. Structured peer review questions

The list of questions (as they were presented to the reviewers) is presented below including information on *Comments-to-Author* and *Comments-to-Editor* fields and text that followed. At the time of the pilot study the questions were also made available on the [Elsevier structured peer review question bank website](https://www.elsevier.com/reviewers/how-to-review/structured-peer-review), and their implementation and the way reviewers fill them out is presented in an [online video](https://www.youtube.com/watch?v=n5K0h8KUtDk). Following this pilot and results, the questions on that website were updated to include our current recommendations.

The reviewers of the pilot journals were informed about these question within the reviewer invitation and the follow up reviewer instruction letter that was set to those who agreed to review a manuscript. The questions were added right above the field of ‘reviewer comment to the author’ and were introduced first with the below introduction:

--------------

Note: In order to effectively convey your recommendations for improvement to the author(s), and help editors make well-informed and efficient decisions, we ask you to answer the following specific questions about the manuscript and provide additional suggestions where appropriate.

1. Are the objectives and the rationale of the study clearly stated?

Please provide suggestions to the author(s) on how to improve the clarity of the objectives and rationale of the study. Please number each suggestion so that author(s) can more easily respond.

2. If applicable, is the application/theory/method/study reported in sufficient detail to allow for its replicability and/or reproducibility?

Please provide suggestions to the author(s) on how to improve the replicability/reproducibility of their study. Please number each suggestion so that the author(s) can more easily respond.

Mark as appropriate with an X:

Yes [] No [] N/A []

Provide further comments here:

3. If applicable, are statistical analyses, controls, sampling mechanism, and statistical reporting (e.g., P-values, CIs, effect sizes) appropriate and well described?

Please clearly indicate if the manuscript requires additional peer review by a statistician.

Kindly provide suggestions to the author(s) on how to improve the statistical analyses, controls, sampling mechanism, or statistical reporting. Please number each suggestion so that the author(s) can more easily respond.

Please number each suggestion so that the author(s) can more easily respond.

Mark as appropriate with an X:

Yes [] No [] N/A []

Provide further comments here:

4. Could the manuscript benefit from additional tables or figures, or from improving or removing (some of the) existing ones? Please provide specific suggestions for improvements, removals, or additions of figures or tables. Please number each suggestion so that author(s) can more easily respond.

5. If applicable, are the interpretation of results and study conclusions supported by the data? Please provide suggestions (if needed) to the author(s) on how to improve, tone down, or expand the study interpretations/conclusions. Please number each suggestion so that the author(s) can more easily respond.

Mark as appropriate with an X:

Yes [] No [] N/A []

Provide further comments here:

6. Have the authors clearly emphasized the strengths of their study/theory/methods/argument?

Please provide suggestions to the author(s) on how to better emphasize the strengths of their study. Please number each suggestion so that the author(s) can more easily respond.

7. Have the authors clearly stated the limitations of their study/methods?

Please list the limitations that the author(s) need to add or emphasize. Please number each limitation so that author(s) can more easily respond.

8. Does the manuscript structure, flow or writing need improving (e.g., the addition of subheadings, shortening of text, reorganization of sections, or moving details from one section to another)?

Please provide suggestions to the author(s) on how to improve the manuscript structure and flow. Please number each suggestion so that author(s) can more easily respond.

9. Could the manuscript benefit from language editing?

***Comments-to-Author***

This field is optional. If you have any additional suggestions beyond those relevant to the questions above, please number and list them here.

***Comments-to-Editor***

This field is optional. If you have any confidential notes for the editor beyond those relevant to the questions above, please number and list them here.

----

Note: The texts “The field is optional…” in the above two sections, was introduced during the pilot. Previously these were just empty fields.

### Appendix Box 2. Word count formula we used in MS Excel

For text in cell A2, the formula was “=LEN(TRIM(A2))-LEN(SUBSTITUTE(A2," ",""))+1”

The same formula was applied for all cells with review answers.

### Appendix Table 2. Reviewer recommendations

Journals use different categories of recommendations. Below is the distribution based on exact wording used by the journals. Absolute Agreement with these terms was 41%.

| **Recommendation** | **N** | **%** |
| --- | --- | --- |
| Accept | 14 | 7% |
| Accept with minor changes | 3 | 1% |
| Major Revision | 2 | 1% |
| Major Revision and Re-review | 8 | 4% |
| Major Revisions | 70 | 33% |
| Minor Revision | 2 | 1% |
| Minor Revisions | 44 | 21% |
| Moderate Revision | 5 | 2% |
| Reconsider after Major Revision | 6 | 3% |
| Reconsider after Minor Revision | 3 | 1% |
| Reject | 50 | 23% |
| Resubmission after major revision | 5 | 2% |
| Revision necessary | 2 | 1% |
| **Total** | 214 | 100.0% |

When the above recommendations were re-coded to accept (accept and accept with minor revisions), reject, and revise (all other categories), agreement rose to 60%.

### Appendix Table 3. Coder inter-rater agreement

Based on answers to 1712 questions.

| Agreement | 1614 (94%) |
| --- | --- |
| Disagreement | 98 (6%) |

### Appendix Self-Review Section

1. Are the objectives and the rationale of the study clearly stated?

Yes, the introduction introduces the reasons why structured peer review may be seen as an improvement to the traditional peer review approach, and the objectives were ordered in the way they were devised, analysed, and presented in the manuscript.

2. If applicable, is the application/theory/method/study reported in sufficient detail to allow for its replicability and/or reproducibility?

Partially, review answers cannot be publicly shared, and so rater coding cannot be replicated without access to that data and signing a non-disclosure agreement. Nevertheless, all codes, and an anonymised dataset is shared, alongside the statistical outputs allowing for reproducibility of all other results.

3. If applicable, are statistical analyses, controls, sampling mechanism, and statistical reporting (e.g., P-values, CIs, effect sizes) appropriate and well described?

Please clearly indicate if the manuscript requires additional peer review by a statistician.

Yes, we described all statistical methods in the order they were conducted first in methods, and then followed SAMPL guidelines for their reporting in results.

4. Could the manuscript benefit from additional tables or figures, or from improving or removing (some of the) existing ones?

We leave this decision to the editor, to help readability we only included 1 table and 1 box in the paper, and all other details in the appendix (i.e. additional 6 tables, and 4 text boxes).

5. If applicable, are the interpretation of results and study conclusions supported by the data?

Yes, we clearly stated all interpretations that come from the data, and called for additional studies for questions that may naturally rise, but were not covered in this study.

6. Have the authors clearly emphasized the strengths of their study/theory/methods/argument?

Yes, we have emphasised that this is the first pilot study of structured peer review, and that it was conducted using random sampling of journals across all disciplines and IF quartiles.

7. Have the authors clearly stated the limitations of their study/methods?

Yes, we have a whole paragraph of limitations in the discussion section.

8. Does the manuscript structure, flow or writing need improving (e.g., the addition of subheadings, shortening of text, reorganization of sections, or moving details from one section to another)?

We leave this decision to the editor, in our view the structure is appropriate.

9. Could the manuscript benefit from language editing?

We have done our best to language proof the manuscript, and we leave the final judgment for this and other questions to the editors and peer reviewers.

### Appendix Table 4. Answer Styles of Reviewers

| **Style** | **n (%)** |
| --- | --- |
| Skipped (i.e. left an answer field blank) for at least one question | 18 (9%) |
| Answered at least one question with see attachment | 16 (7%) |
| Answered at least one question with see comments to authors or answer to other questions | 81 (38%) |
| Did not answer the question in its designated field (i.e., wrote text that was either a review or a comment, not related to the question asked) | 18 (9%) |

### Appendix Table 5. Coded Answers per Question

| **Answer (code)** | **Q1** | **Q2** | **Q3** | **Q4** | **Q5** | **Q6** | **Q7** | **Q8** | **Q9** |
| --- | --- | --- | --- | --- | --- | --- | --- | --- | --- |
| Yes (1) | 104 (49) | 87 (41) | 71 (33) | 13 (6) | 86 (40) | 95 (44) | 73 (34) | 29 (14) | 87 (41) |
| No (2) | 6 (3) | 16 (7) | 16 (7) | 69 (32) | 10 (5) | 26 (12) | 42 (20) | 72 (34) | 127 (59) |
| Not Applicable ( )3 | 4 (2) | 5 (2) | 49 (23) | 10 (5) | 13 (6) | 5 (2) | 7 (3) | 7 (3) | NA |
| Yes, but here are my suggestions for improvement… (4) | 38 (18) | 27 (13) | 15 (7) | 17 (8) | 36 (17) | 23 (11) | 12 (6) | 39 (18) | NA |
| No, but here are my suggestions for improvement… (5) | 32 (15) | 64 (30) | 45 (21) | 8 (4) | 44 (21) | 11 (5) | 27 (13) | 46 (21) | NA |
| NA, but here are my suggestions for improvement… (6) | 0 (0) | 1 (0) | 0 (0) | 0 (0) | 0 (0) | 0 (0) | 0 (0) | 0 (0) | NA |
| Here are my suggestions for improvement… (without specifying yes or no) (7) | 13 (6) | 4 (2) | 6 (3) | 78 (36) | 11 (5) | 28 (13) | 41 (19) | 14 (7) | NA |
| No answer (8) | 0 (0) | 6 (3) | 5 (2) | 1 (0) | **10 (5)** | 1 (0) | 0 (0) | 1 (0) | NA |
| See attached* (9) | 5 (2) | 0 (0) | 2 (1) | 6 (3) | 1 (0) | 5 (2) | 5 (2) | 5 (2) | NA |
| Comments not answering the question (12) | 9 (4) | 1 (0) | 0 (0) | 12 (6) | 0 (0) | 17 (8) | 7 (3) | 0 (0) | NA |
| Yes, but see (14) attached* | 3 (1) | 2 (1) | 1 (0) | 0 (0) | 1 (0) | 0 (0) | 0 (0) | 0 (0) | NA |
| No, but see (15) attached* | 0 (0) | 0 (0) | 1 (0) | 0 (0) | 1 (0) | 1 (0) | 0 (0) | 1 (0) | NA |
| Reviewer unable to assess the question (16) | 0 (0) | 1 (0) | 3 (1) | 0 (0) | 1 (0) | 2 (1) | 0 (0) | 0 (0) | NA |
| Words (Md, IQR)** | 15 (1 to 53) | 27 (11 to 68) | 17 (6 to 71) | 9 (1 to 25) | 22 (9 to 94) | 7 (1 to 19) | 6 (1 to 24) | 8 (2 to 27) | NA |

*Attached documents were not available in this study for analysis.

** Number of words across all questions was Md=161 words (IQR 73 to 157)

### Appendix Table 6. Detailed Agreement per question

| **Question** | **Absolute Agreement** | **Partial Agreement** | **Disagreement** | **Unable to Assess** |
| --- | --- | --- | --- | --- |
| Q1. Are the objectives and the rationale of the study clearly stated? | 36% | 22% | 25% | 17% |
| Q2. If applicable, is the application/theory/method/study reported in sufficient detail to allow for its replicability and/or reproducibility? | 45% | 15% | 31% | 9% |
| Q3. If applicable, are statistical analyses, controls, sampling mechanism, and statistical reporting (e.g., P-values, CIs, effect sizes) appropriate and well described? | 35% | 18% | 23% | 24% |
| Q4. Could the manuscript benefit from additional tables or figures, or from improving or removing (some of the) existing ones? | 25% | 39% | 12% | 23% |
| Q5. If applicable, are the interpretation of results and study conclusions supported by the data? | 32% | 21% | 28% | 19% |
| Q6. Have the authors clearly emphasized the strengths of their study/theory/methods/argument? | 36% | 27% | 13% | 23% |
| Q7. Have the authors clearly stated the limitations of their study/methods? | 24% | 43% | 16% | 17% |
| Q8. Does the manuscript structure, flow or writing need improving (e.g., the addition of subheadings, shortening of text, reorganization of sections, or moving details from one section to another)? | 34% | 38% | 17% | 11% |
| Q9. Could the manuscript benefit from language editing?* | 58% | NA | 42% | NA |

*All questions had open text fields, except question 9 which only had a drop down menu with options yes or no.

### Appendix Box 3. Answer Examples

**Q1**

Examples of reviewers responding *Yes* to question: *Are the objectives and the rationale of the study clearly stated?* - using 1 word, 11 and 44 words are shown below:

*“Yes.”*

*“The objectives and the rationale of the study is clearly stated.”*

*“The objectives of the study to unpick enablers and barriers to equal participation are very clear and are well su*bstantiate*d in the literature. The justification for the research is clearly laid out and describes an important and in-depth examination of girls' ventures into skateboarding.”*

**Q2**

We observed two authors commenting on the appropriateness of the question in regards to the manuscript: “Since the study is based on qualitative interviews, the criteria of replicability or reproducibility do not apply in a strict sense.", and: “It is possible to replicate the study but the is no need to do it. The study design is not appropriate”, with one reviewer stating inability to answer the question: *“The manuscript presented valuable data. However, it is hard to follow the main points and conclusions thus difficult to assess the replicability/reproducibility.”* A detailed yes answer is also presented for comparison: *“Yes, the methods and experiments are described in detail, and all approaches being compared have publicly available code implementations. The appendix even provides the exact parameter values used in the experiments.”*

**Q3**

As this question also recommended reviewers *“clearly indicate if the manuscript requires additional peer review by a statistician”*, we observed that 3(1%) did so, with another 1 (0) specifically stating that: *“The manuscript does not require additional peer review by a statistician.”*

**Q5**

We observed an author reflecting that this question will depend on how authors reply to his other remarks:*“I think this question can go from a No to a Yes depending on responses and modifications mentioned above."*

**Q6**

We observed one author stating they *“cannot assess before the manuscript is revised”*, and another that it *“is difficult to ascertain at present given general method and analyses weaknesses.”*

**Q7**

We observed one author stating limitations are “*not an issue*”, and a few stating authors should list them, but not stating which ones, and one stating *“No, the authors have not stated the limitations. However, it was acceptable since they finished the analysis.”*

**Q8**

We observed that several authors commented on the quality of the English language, which was asked of them again in question 9 (we therefore revised this question to remove the “writing” part and have the language and writing be commented only in question 9 - see Appendix Box 3).

### Appendix Box 4. Recommendations for refining the initial set of 9 questions

Q1. Are the primary (and secondary) objectives clearly stated at the end of the introduction?

[ ] Not Applicable

[ ] Beyond my expertise, additional reviewer(s) should be consulted

[ ] Yes

[ ] No, the authors should (consider): (please list and number Your suggestions so that the author(s) can more easily follow your instructions or provide rebuttals)

Q2. Are the study methods (including theory/applicability/modelling) reported in sufficient detail to allow for their replicability or reproducibility?

[ ] Not Applicable

[ ] Beyond my expertise, additional reviewer(s) should be consulted

[ ] Yes

[ ] No, the authors should (consider): (please list and number Your suggestions so that the author(s) can more easily follow your instructions or provide rebuttals)

Q3. Are statistical analyses, controls, sampling mechanism, and statistical reporting (e.g., P-values, CIs, effect sizes) appropriate and well described?

[ ] Not Applicable

[ ] Beyond my expertise, additional reviewer(s) should be consulted

[ ] Yes

[ ] No, the authors should (consider): (please list and number Your suggestions so that the author(s) can more easily follow your instructions or provide rebuttals)

Q4. Is the clarity and the number of tables and figures in the manuscript appropriate?

[ ] Not Applicable

[ ] Beyond my expertise, additional reviewer(s) should be consulted

[ ] Yes

[ ] No, the authors should (consider): (please list and number Your suggestions so that the author(s) can more easily follow your instructions or provide rebuttals)

Q5. Are the interpretation of results and study conclusions supported by the data and the study design?

[ ] Not Applicable

[ ] Beyond my expertise, additional reviewer(s) should be consulted

[ ] Yes

[ ] No, the authors should (consider): (please list and number Your suggestions so that the author(s) can more easily follow your instructions or provide rebuttals)

Q6. In their discussion section, have the authors clearly emphasized the strengths of their study/theory/methods/argument?

[ ] Not Applicable

[ ] Beyond my expertise, additional reviewer(s) should be consulted

[ ] Yes

[ ] No, the authors should (consider): (please list and number Your suggestions so that the author(s) can more easily follow your instructions or provide rebuttals)

Q7. In their discussion section, have the authors clearly emphasized the limitations of their study/theory/methods/argument?

[ ] Not Applicable

[ ] Beyond my expertise, additional reviewer(s) should be consulted

[ ] Yes

[ ] No, the authors should (consider): (please list and number Your suggestions so that the author(s) can more easily follow your instructions or provide rebuttals)

Q8. Is the manuscript structure and flow clear (e.g., is the manuscript concise, with no text redundancies, and with appropriate subheadings and their order)?

[ ] Not Applicable

[ ] Beyond my expertise, additional reviewer(s) should be consulted

[ ] Yes

[ ] No, the authors should (consider): (please list and number Your suggestions so that the author(s) can more easily follow your instructions or provide rebuttals)

Q9. Is the language of the manuscript appropriate (i.e. without need for language editing)?

[ ] Not Applicable

[ ] Beyond my expertise, additional reviewer(s) should be consulted

[ ] Yes

[ ] No, the authors should (consider): (please list and number Your suggestions so that the author(s) can more easily follow your instructions or provide rebuttals)
